# Decoupled evolution of antimicrobial repertoires in lichen-forming fungi

**DOI:** 10.64898/2026.09.03.749242

**Authors:** Ciaran Kelly, Edgar L. Y. Wong, Anouk van Westerhoven, Vivien Joisten-Rosenthal, Hanna Rövenich, Björn Usadel, Imke Schmitt, Bart P.H.J. Thomma

**Affiliations:** Institute for Plant Sciences, CEPLAS, University of Cologne, Cologne; Senckenberg – Leibniz Institution for Biodiversity and Earth System Research, Senckenberg Biodiversity and Climate Research Centre (SBiK-F), Senckenberganlage 25, 60325 Frankfurt am Main, Germany; Goethe University Frankfurt, Department of Biosciences, Institute of Ecology, Evolution & Diversity, Frankfurt, Germany; Heinrich-Heine University Düsseldorf, Faculty of Mathematics and Natural Sciences, CEPLAS, Institute for Biological Data Science, Düsseldorf, Germany

**Keywords:** antimicrobial protein, effector, mycobiont, photobiont, repertoire evolution, microbiome, secretome, symbiosis

## Abstract

Lichens are stable multipartite symbioses in which a fungal mycobiont associates with one or more photosynthetic partners while hosting complex microbiomes. Secreted antimicrobial proteins (AMPs) have recently emerged as important mediators of fungal interactions with surrounding microbial communities, yet their evolutionary distribution and diversity in lichen-forming fungi remain largely unexplored. Here, we predicted AMP repertoires across 80 phylogenetically diverse lichen-forming fungi representing three photobiont association types. Contrary to our expectation, average repertoire size did not differ between chlorolichens, cyanolichens and tripartite lichens (363 AMPs per species overall), even though repertoire size varied almost five-fold among species and 23 AMP families showed photobiont-associated enrichment or depletion. This quantitative conservation masks extensive compositional turnover, with families occurring in at least 90% of species coexisting alongside numerous lineage-specific groups. Pooled population sequencing of 11 *Umbilicaria phaea* populations along two elevational gradients further revealed recurrent AMP presence– absence variation within a single species. In contrast to AMPs, biosynthetic gene clusters (BGCs) and carbohydrate-active enzymes (CAZymes), which include known antimicrobials, were markedly reduced in cyanolichens relative to chlorolichens. Lichen-forming fungi thus combine a conserved baseline antimicrobial protein capacity, whose composition nonetheless turns over between lineages and populations, with chemical and enzymatic systems that differentiate along photobiont association and may support ecological and lifestyle-associated adaptation.

**SIGNIFICANCE STATEMENT:** Lichen-forming fungi interact simultaneously with photosynthetic partners and diverse microbial communities, yet how their antimicrobial capacity evolves in this complex symbiotic context is poorly understood. By comparing 80 lichen-forming fungal genomes, we show that the size of their predicted antimicrobial protein repertoires is broadly conserved across associations with green algae, cyanobacteria, or both, despite extensive turnover in the constituent protein families across lineages and populations. In contrast, carbohydrate-active enzymes and biosynthetic gene clusters vary substantially with photobiont association. These findings reveal a decoupling of antimicrobial repertoire size from repertoire composition and from other antimicrobial molecules, suggesting that distinct molecular repertoires are subject to different evolutionary constraints. Thus, lichen-forming fungi maintain a broadly conserved genomic potential for protein-based antimicrobial activity while retaining substantial molecular flexibility.

## INTRODUCTION

Lichens are stable symbioses in which a fungus, termed the mycobiont, associates with a photosynthetic partner, termed the photobiont, to form a structured thallus (Hale, 1974; Henssen and Jahns, 1974; Ahmadjian, 1993; Nash, 2008; Hawksworth and Grube, 2020). Such lichen thalli occur in a wide diversity of appearances, shapes and colours, reflecting the remarkable morphological and ecological diversity of lichens (Hale, 1974; Henssen and Jahns, 1974; Nash, 2008). Lichens occur in nearly all terrestrial ecosystems on Earth (Spribille et al., 2022), including extreme habitats where they often dominate as relatively few other organisms persist (Pichler et al., 2023). While the mycobiont is the namesake of the lichen, the photosynthetic partner is either a green alga, a cyanobacterium or, in particular cases, both (Ahmadjian, 1993; Nash, 2008; Rodríguez-Arribas et al., 2023). Lichens that harbour only green algae, also referred to as (bipartite) chlorolichens, are widespread, while lichens that harbour only cyanobacteria, known as (bipartite) cyanolichens, or both photobiont types, named cephalolichens or tripartite lichens, are comparatively rare among lichen-forming fungi (Honegger, 2008; Nash, 2008; Rodríguez-Arribas et al., 2023). In chlorolichens, green algae serve as sole photobionts and supply the mycobiont with photosynthetically fixed carbon. In contrast, in cyanolichens, a cyanobacterial photobiont provides both fixed carbon through photosynthesis and fixed nitrogen through biological nitrogen fixation. In cephalolichens, however, these functions are partitioned between the two photosynthetic organisms: the green alga is the principal photobiont and source of fixed carbon, while the cyanobacterium, localized in cephalodia, primarily supplies fixed nitrogen (Nash, 2008; Rascio & La Rocca, 2013). Intriguingly, some lichen-forming fungi can switch photobiont lineages, typically acquiring a different species or strain within the same photobiont type, presumably to adapt to changing environments (Škvorová et al., 2022; Kantnerová & Škaloud, 2025). For example, populations of *Umbilicaria phaea* and *U. pustulata* along elevational gradients revealed elevation-associated switching among algal photobionts belonging to different *Trebouxia* lineages (Dal Grande et al., 2018; Rolshausen et al., 2020; Rolshausen et al., 2023).

Beyond the primary symbiosis between mycobiont and photobiont, lichens host diverse communities of associated microorganisms, comprising other fungi that include basidiomycete yeasts (Spribille et al., 2016; Tuovinen et al., 2019) and black fungi (Keller et al., 2025; Keller et al., 2026), as well as bacterial communities (Grube et al., 2009; Tagirdzhanova et al., 2024). These communities have collectively been referred to as the lichen-associated microbiota (Grube et al., 2015). Sequencing studies have shown that lichen-associated microbiota differ between species of lichen-forming fungi and can be vertically transmitted via vegetative propagules (Aschenbrenner et al., 2014; Keller et al., 2026). Large-scale metagenomic analyses have further revealed dominant, recurrent bacterial lineages and conserved symbiont associations across hundreds of samples, providing genomic evidence for integrated lichen microbiomes (Tagirdzhanova et al., 2024). However, while these studies provide a descriptive framework of lichen-associated microbial diversity, the molecular mechanisms that govern the establishment and maintenance of these communities within the lichen thallus remain largely unresolved, particularly with respect to how the mycobiont or photobiont contributes to the assembly and stability of these communities. Currently, it also remains unclear whether these lichen-associated microbiota merely inhabit the lichen as a niche or contribute to the functioning of the lichen as a symbiotic “holobiont”.

As lichens represent evolutionarily ancient symbioses that date back to the Palaeozoic era (Taylor et al., 1995), the mechanisms underlying microbial interactions within the thallus are likely rooted in early microbial interactions. Ancient environments were characterized by dense microbial communities (Noffke et al., 2006; Parfrey et al., 2011), where intense intermicrobial competition for scarce nutrients and habitats likely drove the evolution of diverse competition mechanisms, including a wide diversity of antimicrobial molecules (Snelders et al., 2022). In lichens, the presence of such antimicrobial molecules has been demonstrated using lichen extracts, which exhibit both antibacterial and antifungal activities (e.g. Güllüce et al., 2006). Further studies have identified antimicrobial secondary metabolites produced by lichens (Türk et al., 2006) and comparative genomic analyses have revealed that lichen-forming fungi encode numerous biosynthetic gene clusters associated with diverse metabolites, including compounds with antimicrobial activity (Singh et al., 2025). In addition to specialized metabolites, fungal carbohydrate-active enzymes (CAZymes) can mediate interactions with other organisms by modifying or degrading polysaccharides, including those present in microbial cell walls (Wardman et al., 2022; Zheng et al., 2023). CAZymes are abundant in the genomes of lichen-forming fungi, and show lineage-specific distributions (Resl et al., 2022). Recently, we have demonstrated that fungi, irrespective of their lifestyle, encode extensive repertoires of antimicrobial proteins (AMPs) that play roles in microbial antagonism and niche competition (Mesny et al., 2024; Mesny et al., 2026). Here, we assess the distribution and diversity of AMP catalogues across lichen-forming fungi. Moreover, we investigate whether the association with different photobionts, green algae vs. cyanobacteria, is reflected by differences in mycobiont AMP catalogues. We subsequently extend these analyses to BGC and CAZyme repertoires, given that these also comprise antimicrobials. Together, these complementary repertoires allow us to assemble a comprehensive picture of the antimicrobial potential of lichen-forming fungi across their phylogenetic and symbiotic diversity.

## RESULTS

### Composition of a phylogenetically and genomically diverse mycobiont genome collection

To assess whether mycobionts differ in their antimicrobial molecule catalogues, we assembled a broad dataset of 80 genomes from the fungal phylum Ascomycota, which contains over 98% of lichen-forming fungi (Lutzoni and Miadlikowska, 2009). Of these, 67 genomes belong to the Lecanoromycetes, which harbours the majority of described mycobionts (Miadlikowska et al., 2006). Genome completeness was evaluated using BUSCO benchmarks and ranged from 78.1% to 99.1% (Fig. 1; Table S1). The selection contained 49 genera that are represented by a single species and 11 genera with more than one species. While 70 species are chlorolichens, five are cyanolichens, and another five are tripartite lichens (Fig. 1). This distribution reflects global photobiont proportions, where 85–90% of described species associate with green algae and significantly fewer partner with cyanobacteria or both photobionts (Jung et al., 2024). Thus, our dataset comprises a wide diversity of lichen-forming fungi and reflects the dominance of Lecanoromycetes and chlorolichens among lichen-forming fungi.

**Figure 1.**
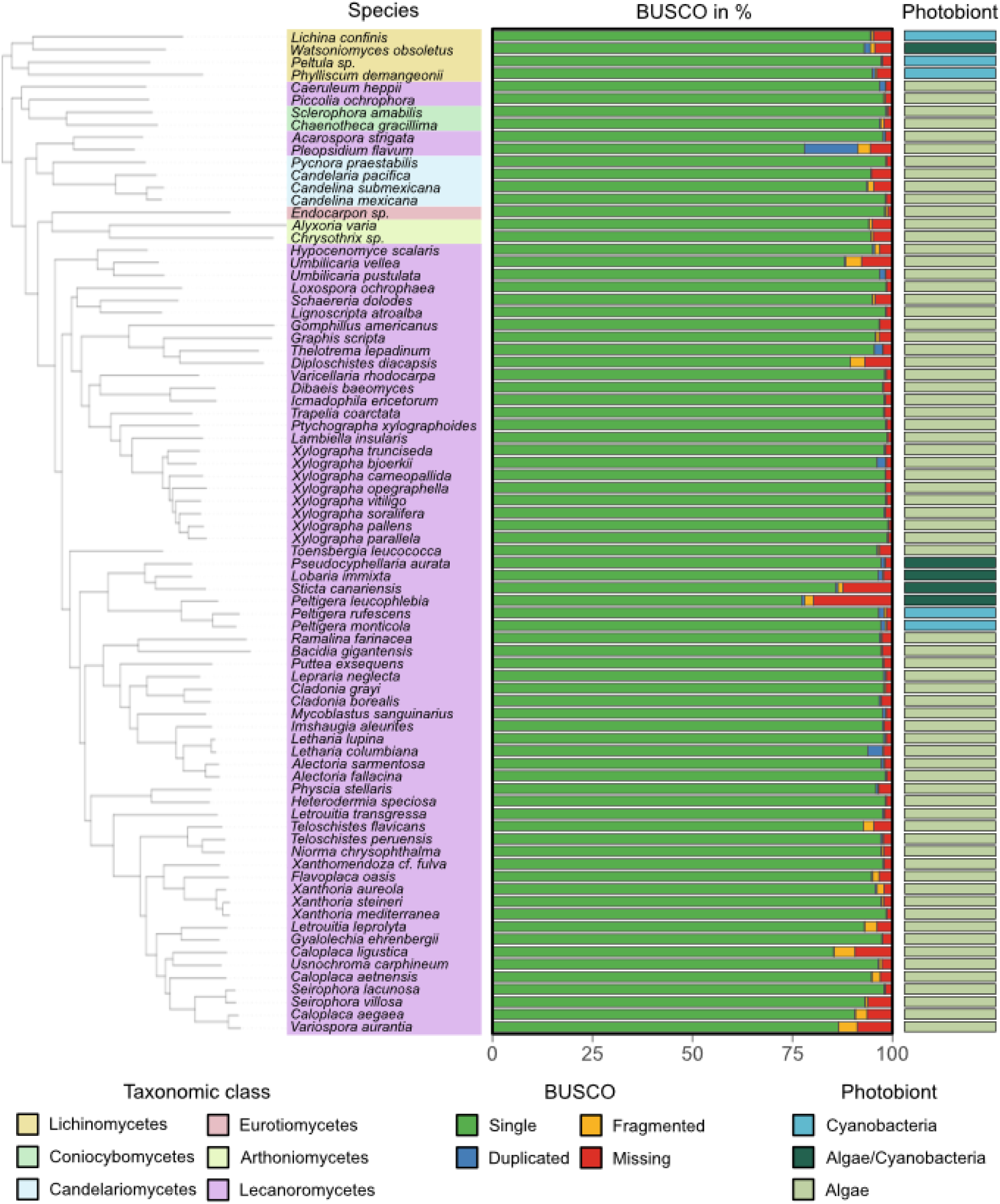
Phylogeny, genome completeness and photobiont associations of 80 lichen-forming fungi. Left panel: Phylogenomic tree based on all protein-coding genes. Species names are colour-coded according to their taxonomic class assignment. Middle panel: Stacked bar plot showing BUSCO statistics (completeness) of single, duplicated, fragmented and missing genes. Right panel: Indication of the associated photobiont for each species.

Next, we generated a phylogenetic tree based on all protein-coding genes and observed two distantly related clades, each containing bipartite cyanolichens and tripartite lichens (Fig. 1). As in previous phylogenies (Diaz-Escandon et al., 2022), one cluster belonged to the Lecanoromycetes, whereas the other was positioned within the Lichinomycetes, suggesting at least two independent evolutionary origins of cyanobacterial lichen symbioses.

### Different photobiont association types encode comparable numbers of antimicrobial proteins

To assess the diversity and distribution of AMP repertoires in lichen-forming fungi, we screened the genomes of the 80 species for genes encoding AMPs using the recently developed AMAPEC software, which predicts antimicrobial activity among secreted effector proteins based on their physicochemical properties (Mesny et al., 2026). On average, mycobionts encoded 363 AMPs per species, although repertoire sizes varied substantially across taxa (Fig. 2A). Total AMP numbers ranged from 138 in *Lichina confinis* to 635 in *Physcia stellaris*. The AMP repertoire size strongly correlated with the total secretome size (Spearman ρ = 0.97), while the correlation with genome size was weaker (Spearman ρ = 0.56) (Fig. S1; Table S1 and S2). While the five bipartite cyanolichens encoded 328 AMPs on average, this average did not differ significantly from the averages found for the 70 bipartite chlorolichens (364) and the five tripartite lichens (383) (Fig. 2A; Table S2).

**Figure 2.**
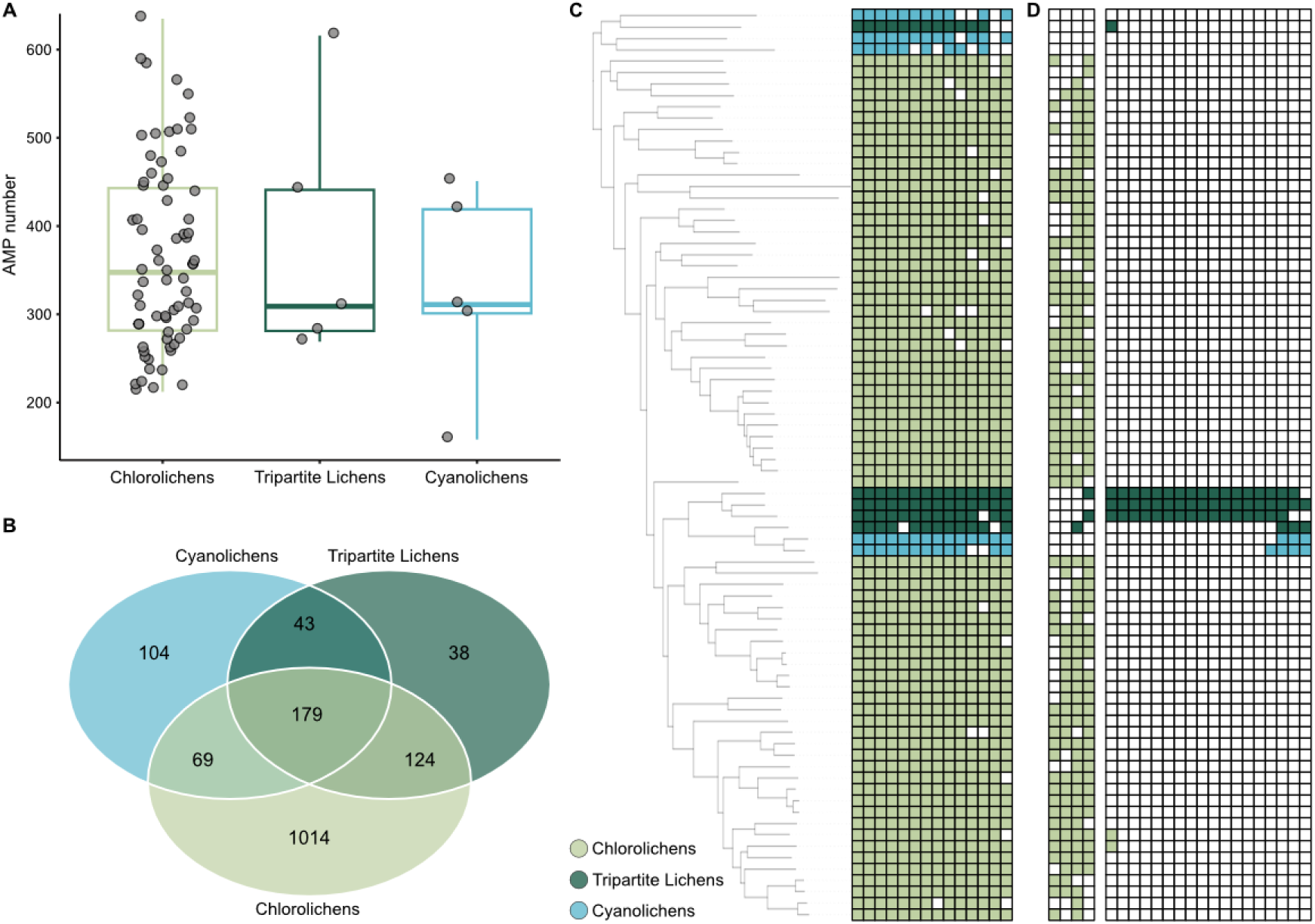
Diversity, conservation and distribution of antimicrobial proteins (AMPs) across 80 genomes of lichenized fungi. **A)** Box plot showing the total number of predicted AMPs encoded by mycobiont genomes grouped according to photobiont association. Each dot represents a single species. No statistically significant differences were detected between photobiont association types. The outlines of the box plots are colour-coded for bipartite cyanolichens (blue), bipartite chlorolichens (pale green), and tripartite lichens (dark green). **B)** Venn diagram showing the distribution of AMP orthogroups among bipartite cyanolichens, bipartite chlorolichens and tripartite lichens. Numbers indicate orthogroups unique to or shared among the different photobiont association types. **C)** Heatmap showing the presence (squares coloured by photobiont association type: bipartite cyanolichens (blue), bipartite chlorolichens (pale green) and tripartite lichens (dark green)) and absence (empty square) of the 14 AMP orthogroups identified in at least 72 of the 80 analysed species. Species are arranged according to the phylogenetic order shown in Fig. 1. **D)** Heatmap showing the occurrence patterns of 23 AMP orthogroups displaying significant enrichment or depletion among photobiont association types (Fisher’s exact test, adjusted p-value < 0.05). Empty squares indicate absence while colour-filled squares indicate the photobiont association type: bipartite cyanolichens (blue), bipartite chlorolichens (pale green) and tripartite lichens (dark green).

To determine the extent to which AMP repertoires are shared among species, we grouped all 29,043 predicted AMPs into orthologous groups based on sequence similarity. In total, 26,316 AMPs (90.6%) were assigned to 1,570 orthogroups, whereas 2,727 AMPs (9.4%) lacked detectable homologues and were therefore classified as singletons. The proportion of AMPs assigned to orthogroups varied considerably among species, ranging from 58.7% in *Watsoniomyces obsoletus* to 99.4% in *Xanthoria steineri*. Species belonging to genera represented by multiple members in the dataset generally exhibited higher orthogroup assignment rates than species belonging to genera represented by a single member, indicating that taxonomic sampling influences the detection of families of homologous AMPs. To identify potential AMP families that are conserved across lichen-forming fungi, we examined orthogroup conservation and identified 14 orthogroups that were present in at least 90% of the species (72 out of 80; Fig. 2C; Table S3). These conserved orthogroups were detected across the phylogeny and occurred in representatives of all three photobiont association strategies. For ten of these 14 orthogroups Pfam annotations could be assigned. These included defensin-like protein, aspartyl protease, peptidase S10, carboxylesterase, pro-kumamolisin activation, ferritin-like, thioredoxin-related, EMP24/GP25L, tyrosinase, arginase and DnaJ domains. Several of these domains are associated with processes relevant to microbial antagonism or cellular defence, including direct antimicrobial activity, extracellular proteolysis, iron homeostasis, redox protection and melanization (Ganz, 2003; Mygind et al., 2005; Deng et al., 2018; Seo et al., 2018; Claus & Decker, 2006; Schaefer et al., 2020).

We next investigated how AMP orthogroups are distributed across photobiont association types. Among the 1,570 identified orthogroups, 1,014 were detected exclusively in chlorolichens, whereas 104 and 38 orthogroups were unique to cyanolichens and tripartite lichens, respectively. In contrast, only 179 orthogroups were shared among all three photobiont association types (Fig. 2B; Table S3). Fisher’s exact tests identified 23 orthogroups with significantly biased occurrence patterns after correction for multiple testing (adjusted p-value <0.05; Fig. 2D; Table S4). Four orthogroups were significantly enriched in chlorolichens, comprising PI-PLC X, EXPB1-like, arylsulfotransferase and peptidase S28 domains. The remaining 19 orthogroups were significantly depleted in chlorolichens and occurred predominantly in cyanolichens and/or tripartite lichens. For six of them, Pfam annotation revealed SIP1, HpcH/HpaI aldolase, divisome-associated membrane protein, AAA+ ATPase lid, HOXB-AS3 peptide and DUF3530 domains (Table S5). Only a single orthogroup (OG0000001) contains clear homologs to a previously characterized fungal AMP, namely a fungistatic protein from *Chaetomium globosum* (Istifadah et al., 2006).

Altogether, our results indicate that while overall AMP repertoire sizes differ greatly between mycobiont species, but are similar between different photobiont association types. Moreover, only few AMP families can be associated with particular photobiont association types.

### Sequence homology determines structural clustering of lichen antimicrobial proteins

Recent studies have shown that many seemingly unrelated fungal effectors can be grouped into structurally conserved families despite limited sequence similarity, revealing evolutionary relationships that remain undetected by conventional sequence-based analyses (Derbyshire & Raffaele, 2023; Seong & Krasileva, 2023). We therefore compared sequence-based orthogroups with predicted structural clusters to assess whether structural relationships provide additional insights into AMP evolution in lichen-forming fungi. Across the 80 analysed genomes, 29,043 predicted AMPs grouped into 12,941 structural clusters using a permissive similarity threshold (TM-score ≥ 0.50) (Table S6). Bipartite chlorolichens encoded an average of 303 structural clusters, bipartite cyanolichens 301, and tripartite lichens 325 structural clusters per genome. Thus, similar as for total AMP numbers, the average number of structural clusters encoded per genome did not differ significantly among photobiont association types (Kruskal–Wallis, p = 0.96; Fig. 3A).

**Figure 3.**
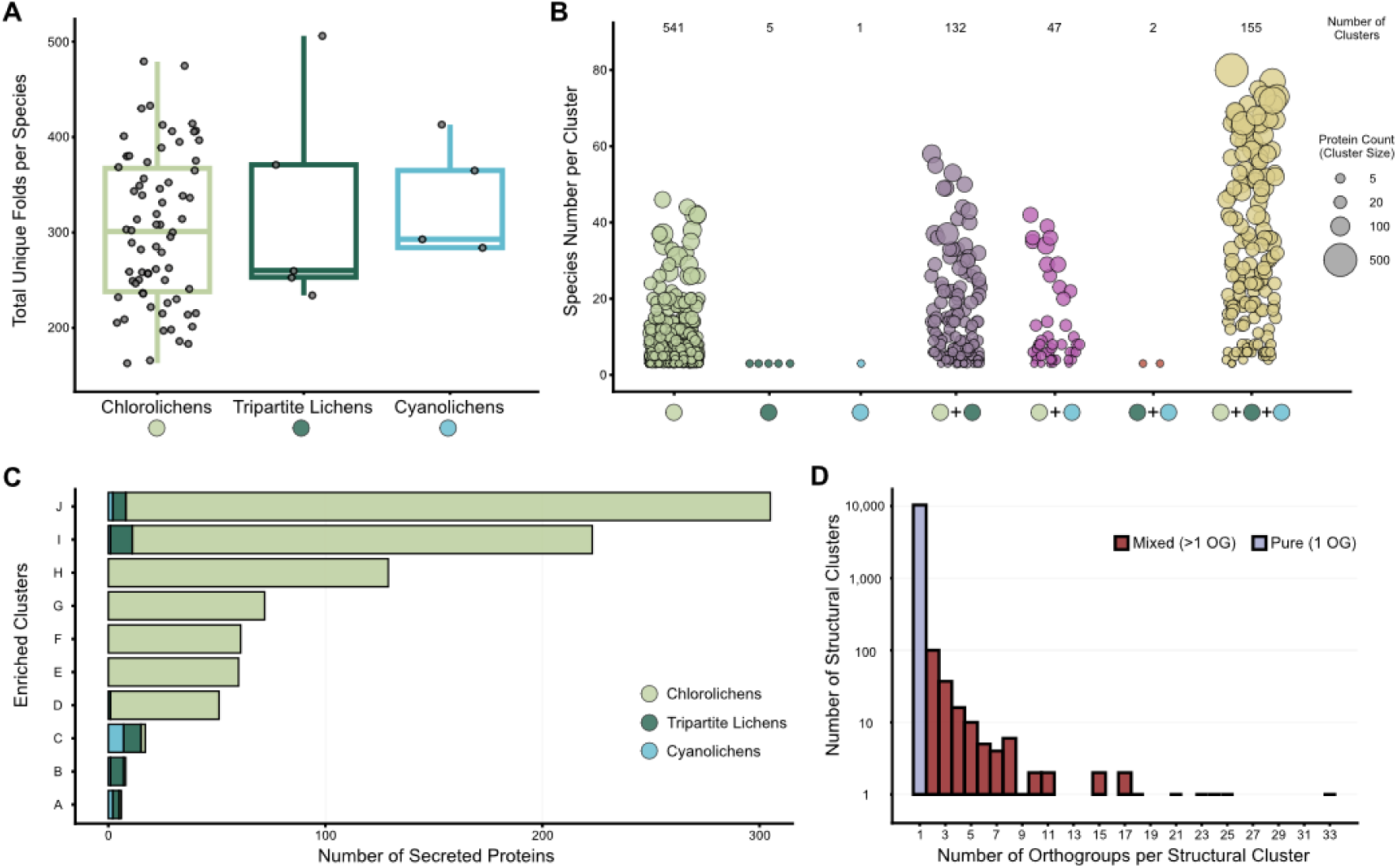
Structural clustering of lichen antimicrobial proteins is driven by sequence homology. **A)** Box plot showing the number of unique AMP structural clusters encoded by mycobiont genomes grouped according to photobiont association. Each dot represents a single species. No statistically significant differences were detected between photobiont association types (Kruskal–Wallis test, p = 0.96). The outlines of the box plots are colour-coded for bipartite cyanolichens (blue), bipartite chlorolichens (pale green), and tripartite lichens (dark green). **B)** Distribution of 883 AMP structural clusters detected in at least three species. Structural clusters are grouped according to their occurrence among bipartite cyanolichens (blue dot), bipartite chlorolichens (pale green dot) and tripartite lichens (dark green dot). Colour coding can also be seen in panel A. Bubble size corresponds to the total number of proteins assigned to each structural cluster. Numbers indicate the total number of structural clusters within each occurrence category. **C)** Bar plot showing the ten AMP structural clusters displaying significant enrichment or depletion among photobiont association types (Fisher’s exact test, adjusted p-value < 0.05). Clusters were labelled from A to J following the number of proteins per cluster. Bars are coloured by photobiont association type: bipartite cyanolichens (blue), bipartite chlorolichens (pale green) and tripartite lichens (dark green). **D)** Bar plot showing the number of sequence-based orthogroups contained within individual structural clusters. Structural clusters containing proteins from a single orthogroup are shown separately from clusters containing proteins from multiple orthogroups.

To evaluate the relationship between structural and sequence-level classification, we compared structural clusters with orthogroups. Structural and sequence groupings showed a high degree of concordance (Normalised Mutual Information = 0.7608). Of the 12,941 structural clusters identified, 10,354 could be assigned to a single orthogroup, whereas 193 (1.83%) clusters contained proteins assigned to multiple orthogroups (Fig. 3D). The remaining 2,394 clusters lacked sufficient sequence similarity for orthogroup assignment. These results indicate that AMP structural clusters are largely determined by sequence homology.

To assess whether AMP structural families show photobiont-associated distributions, we examined how structural clusters are shared across photobiont association types. Among the 883 structural clusters present in at least three species, 541 were restricted to bipartite chlorolichens, whereas 155 were shared across all photobiont association types. In contrast, only eight clusters were detected exclusively in bipartite cyanolichens and/or tripartite lichens (Fig. 3B). Fisher’s exact tests identified ten structural clusters enriched in certain lichen types after correction for multiple testing (adjusted p-value < 0.05; Fig. 3C; Table S4). Seven of the ten clusters contained Pfam-annotated proteins. Five of the seven annotated structural clusters (E, F, G, I and J) overlapped with enriched or conserved AMP orthogroups, whereas clusters A and D did not. Cluster D contained proteins with domains associated with the Ca²⁺ regulator and membrane fusion protein Fig1 and the serpentine-type 7TM GPCR chemoreceptor Srh, whereas cluster A contained proteins with von Willebrand factor type A domains (Table S5). The remaining three enriched structural clusters, B, C and H, lacked Pfam annotations. No homology to previously characterized AMPs was found for proteins from cluster A, B, C, D or H. Together, these results indicate that structural clustering largely recapitulated sequence-based AMP groupings, while also identifying a small number of structurally defined clusters with previously unclassified sequence relationships.

### Intraspecific variation of antimicrobial proteins in *Umbilicaria phaea*

Although the variation in AMP catalogues between photobiont association types is limited, considerable variation exists among species. We therefore assessed variation within a single lichen species. To this end, we analysed previously generated pooled sequencing data from 11 *Umbilicaria phaea* populations, comprised of 50 individuals each, sampled along two elevational gradients in California, USA, one in the Sierra Nevada and the other at Mount San Jacinto (Rolshausen et al., 2020). Photobiont turnover has previously been documented across these elevational gradients, with dominant algal photobionts belonging to different *Trebouxia* species replacing one another at distinct points along the gradients (Rolshausen et al., 2020). A pronounced transition in *Trebouxia* species composition occurs between population 2 (1014 m) and population 3 (1533 m) along the Sierra Nevada gradient, and between population 4 (1324 m) and population 5 (1992 m) along the San Jacinto gradient (Rolshausen et al., 2020). Two reference genomes of *U. phaea* were also available, originating from individuals sampled at the lowest (631 m) and highest (2036 m) sites of the Sierra Nevada gradient (Rolshausen et al., 2020), yielding 246 and 287 predicted AMPs, respectively. Of these, 208 AMP candidates were shared between the two reference genomes, while 38 were unique to the low-elevation reference and 79 to the high-elevation reference. Together, these comprised 325 non-redundant AMPs, pointing to considerable intraspecific variation in AMP catalogues.

Next, we assessed the presence of the 325 non-redundant AMP candidates across the 11 populations. AMP candidates that were present in all populations, restricted to a single population only, or absent in a single population only, were excluded, resulting in 48 candidate AMPs that showed substantial presence–absence variation, and that could be grouped into eight clusters based on their distribution patterns. Two clusters showed particularly clear and recurring patterns across the two elevational gradients; cluster 2 contained four AMPs that were generally absent at lower-elevation sites and present at sites above 1324 m, whereas cluster 8 contained five AMPs that were generally present at lower-elevation sites but absent at sites above 1533 m (Fig. 4; Table S7). Intriguingly, these distribution patterns broadly coincided with the previously described transitions in *Trebouxia* community composition between populations 2 and 3 in the Sierra Nevada and populations 4 and 5 at Mount San Jacinto (Rolshausen et al., 2020). Homology searches against the 80 lichen-forming fungal genomes analysed in this study identified sequence homologs for all nine AMPs. For eight of them, the strongest hit was found in *Umbilicaria vellea* or *Hypocenomyce scalaris*, both belonging to the order Umbilicariales. Seven of the nine proteins belonged to the same orthogroup; OG0000018 (Table S4), containing proteins from 30 mycobionts, all classified as associating exclusively with green algal photobionts but lacking Pfam domain annotation. Together, these results reveal substantial intraspecific variation in AMP catalogues in *U. phaea*.

**Figure 4.**
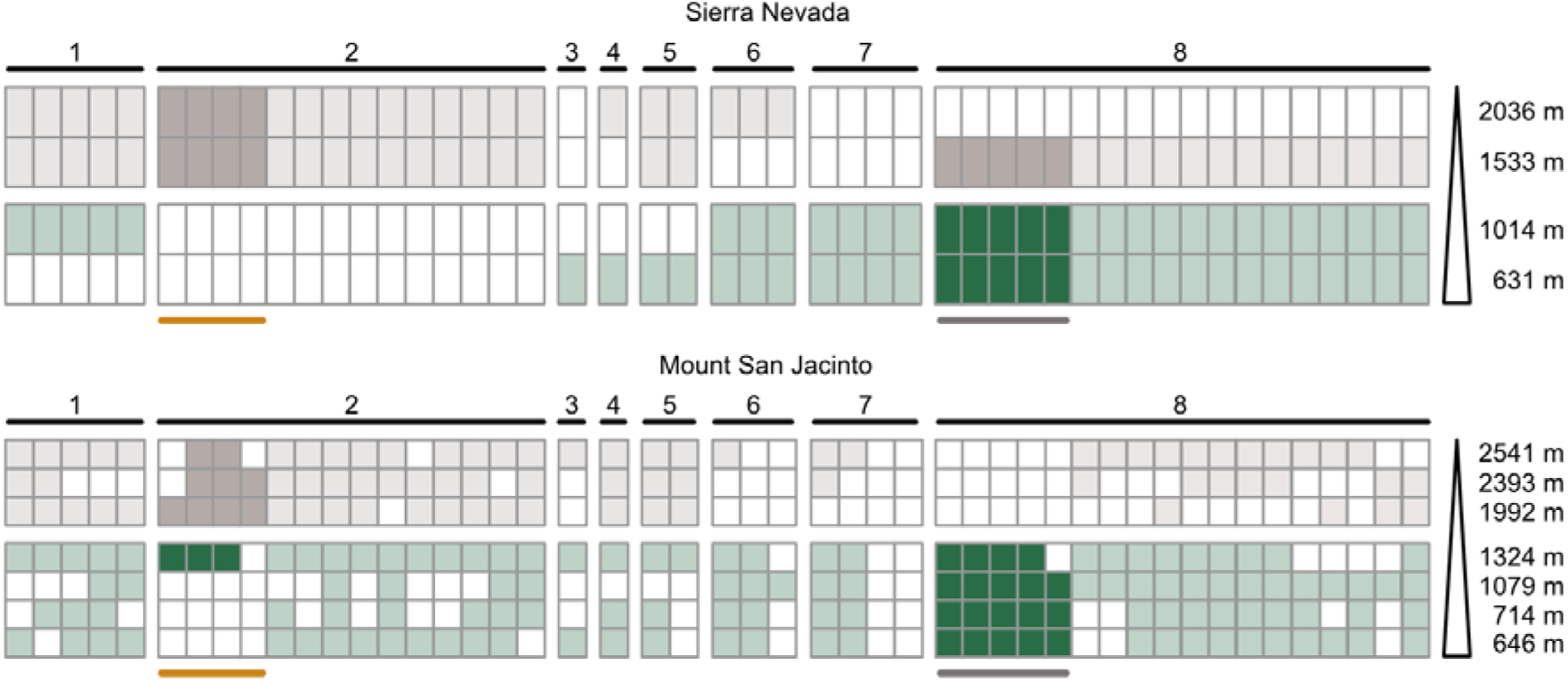
Elevation-associated AMP repertoires in *Umbilicaria phaea*. Presence–absence patterns of 48 AMP-encoding genes showing variation across sampling sites along the Sierra Nevada and Mount San Jacinto gradients. AMPs were clustered according to shared distribution patterns across sampling sites. Samples are coloured according to the previously described *Trebouxia* community transition, with green representing sampling pools below the transition zone and grey representing those above the transition. AMPs showing consistent distribution patterns across the two elevational gradients are highlighted in a darker shade and with a line below. These patterns broadly coincide with the transitions in dominant *Trebouxia* haplotypes described by Rolshausen et al. (2020). The orange and grey lines indicate the dominant *Trebouxia* haplotypes associated with the respective sampling populations, following the colour coding of Rolshausen et al. (2020).

### Cyanolichens encode fewer carbohydrate-active enzymes

Carbohydrate-active enzymes (CAZymes) can display antimicrobial activity by targeting carbohydrates in microbial cell walls (Wardman et al., 2022). We therefore characterized the secreted CAZyme repertoires across 80 lichen-forming fungi and identified an average of 93 CAZymes per species. Total CAZyme numbers ranged from 38 in *Lichina confinis* to 287 in *Thelotrema lepadinum*. As previously observed (Resl et al., 2022), cyanolichens encoded significantly fewer CAZymes than chlorolichens (Dunn’s test, **p < 0.01; Fig. 5A). The five cyanolichen species encoded an average of 56 CAZymes, whereas the 70 chlorolichen species encoded an average of 96 CAZymes. The remaining five tripartite lichens encoded an intermediate average of 78 CAZymes per species (Fig. 5A; Table S8).

**Figure 5.**
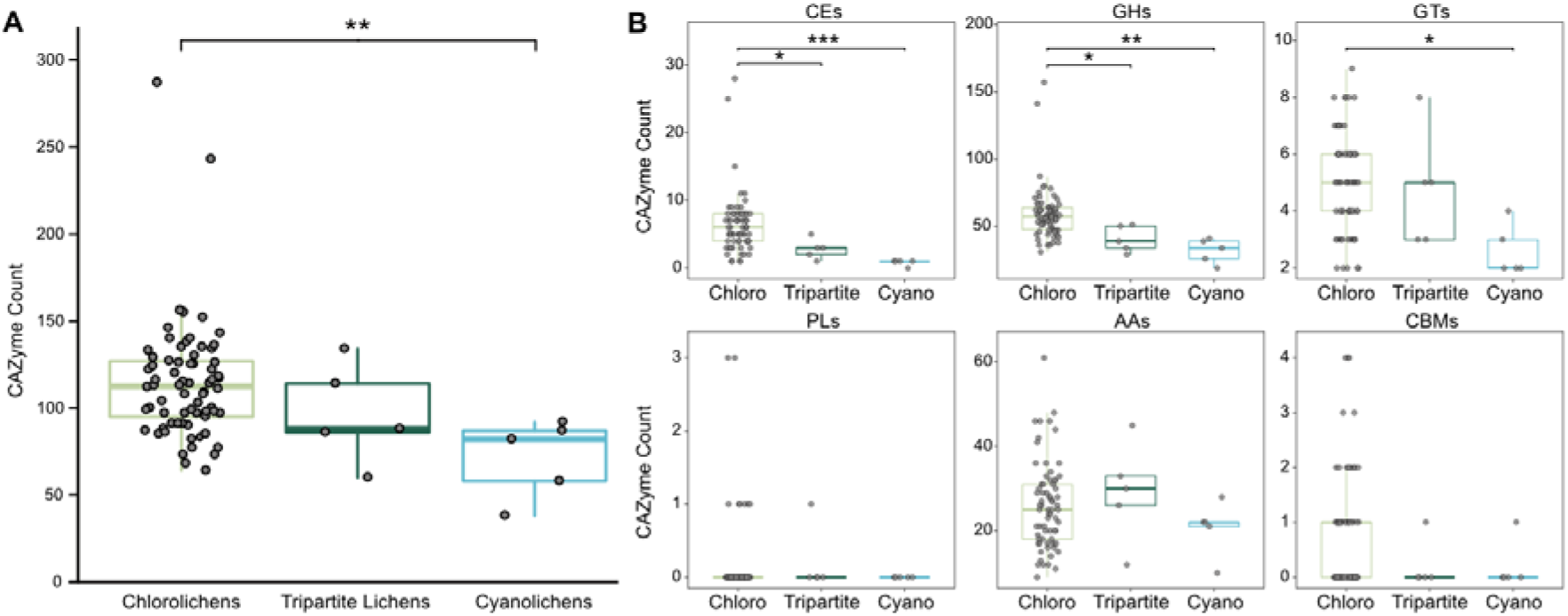
Comparison of CAZyme repertoires across lichen-forming fungi with different photobiont associations. **A)** Box plot showing total numbers of CAZymes encoded by mycobiont genomes grouped according to photobiont association. Bipartite chlorolichens encode significantly more CAZymes than cyanolichens (Dunn’s test, **p < 0.01). The outlines of the box plots are colour-coded for bipartite cyanolichens (blue), bipartite chlorolichens (pale green), and tripartite lichens (dark green). **B)** Box plots showing abundance of individual CAZyme classes including carbohydrate esterases (CEs), glycoside hydrolases (GHs), glycosyltransferases (GTs), polysaccharide lyases (PLs), auxiliary activity enzymes (AAs) and carbohydrate-binding modules (CBMs). Significant differences were detected for CEs, GHs, and GTs between photobiont association groups (Dunn’s test; *p < 0.05, **p < 0.01, ***p < 0.001). The outlines of the box plots are colour-coded for bipartite cyanolichens (blue), bipartite chlorolichens (pale green), and tripartite lichens (dark green). Detailed counts for each CAZyme subclass are provided in Table S8.

We further assessed whether differences in specific CAZyme classes contributed to the reduced repertoires of cyanolichens compared with chlorolichens. We compared the abundance of glycoside hydrolases (GHs), auxiliary activity enzymes (AAs), carbohydrate esterases (CEs), glycosyltransferases (GTs), polysaccharide lyases (PLs) and carbohydrate-binding modules (CBMs) (Drula et al., 2022; Table S8). The strongest difference was observed for CEs (Dunn’s test, ***p < 0.001; Fig. 5B). Cyanolichens encoded on average one CE per species, whereas chlorolichens encoded on average six. Tripartite lichens encoded an intermediate average of three CEs, which remained significantly lower than chlorolichens (Dunn’s test, *p < 0.05; Fig. 5B). Significant differences were also detected for GHs. Chlorolichens encoded on average 58 GHs per species compared to 32 in cyanolichens (Dunn’s test, **p < 0.01; Fig. 5B). Tripartite lichens encoded an average of 41 GHs, which was also significantly lower than chlorolichens (Dunn’s test, *p < 0.05; Fig. 5B). GT abundance likewise differed significantly between chlorolichens and cyanolichens (Dunn’s test, *p < 0.05; Fig. 5B), with chlorolichens encoding on average five GTs per species compared to three in cyanolichens. No significant difference was observed between chlorolichens and tripartite lichens in any CAZyme class. Furthermore, no significant differences were detected for AAs, CBMs, or PLs between any pair of photobiont association types. AA abundance was similar across groups, with averages of 26, 21 and 29 AAs per species in chlorolichens, cyanolichens and tripartite lichens, respectively. CBMs were rare overall and were detected in only 38 of the 80 species, with fewer than one CBM per species on average across all three photobiont association types. PLs were the rarest CAZyme class, occurring in only nine species, and none of the cyanolichens encoded a PL (Table S8).

We next examined CAZyme families with reported activity against microbial or algal cell-wall components. These included families targeting bacterial peptidoglycan (GH22, GH23, GH24, GH25, GH73, GH102, GH103 and GH104), chitin in fungal and algal cell walls (GH18, GH19 and GH20), and β-glucans in fungal and algal cell walls (GH5, GH16, GH17, GH55, GH64, GH81, GH128 and GH152). We also included AA11, which comprises fungal lytic polysaccharide monooxygenases with activity towards chitin (Hemsworth et al., 2014; Støpamo et al., 2021). Collectively, these cell wall-associated CAZyme families were less abundant in cyanolichens, which encoded an average of 19 enzymes per species compared with 31 in chlorolichens (pairwise Wilcoxon test, adjusted *p-value = 0.011; Fig. S2A; Table S8). The strongest differences were observed among the β-glucan-active families GH17 and GH5, targeting β-1,3-glucans and mixed-linkage β-glucans, and mixed-linkage β-1,3-1,4-glucans, respectively, both of which were significantly less abundant in cyanolichens than in chlorolichens (pairwise Wilcoxon test, adjusted **p-values = 0.0087 and 0.0022, respectively; Fig. S2B). Thus, the reduced CAZyme repertoires of cyanolichens also extended to families with reported activity against cell wall components, including peptidoglycan, chitin and β-glucans. Together, these results reveal an overall reduction in CAZyme repertoires in cyanolichens compared with chlorolichens, particularly reflecting lower abundances of CEs, GHs and GTs.

### Cyanolichens encode fewer biosynthetic gene clusters

Fungal secondary metabolites have been implicated in various physiological processes, including microbial antagonism, given that they frequently exhibit antimicrobial properties (Rangel et al., 2021). To assess whether secondary metabolite catalogues differ among lichen-forming fungi, we annotated biosynthetic gene clusters (BGCs) and compared their profiles across the 80 species. On average, the 80 lichen-forming fungi encoded 42 BGCs. Similar as for the CAZymes, but in contrast to the AMP catalogues, cyanolichens contained significantly fewer BGCs than chlorolichens (t-test, *p < 0.05; Fig. 6A) with the five cyanolichens encoding an average of 25 BGCs compared with 44 in the 70 chlorolichens. The remaining five tripartite lichens encoded 40 BGCs on average, although this group includes *Lobaria immixta* that encodes the highest number of BGCs (84) within our dataset (Fig. 6A). The number of BGCs strongly correlated with the number of CAZymes across the 80 species (Spearman ρ = 0.93; Fig. S1A). Together, these patterns indicate that the two repertoires show similar broad differentiation among photobiont association types, with cyanolichens generally encoding fewer BGCs and CAZymes than chlorolichens (Table S9).

**Figure 6.**
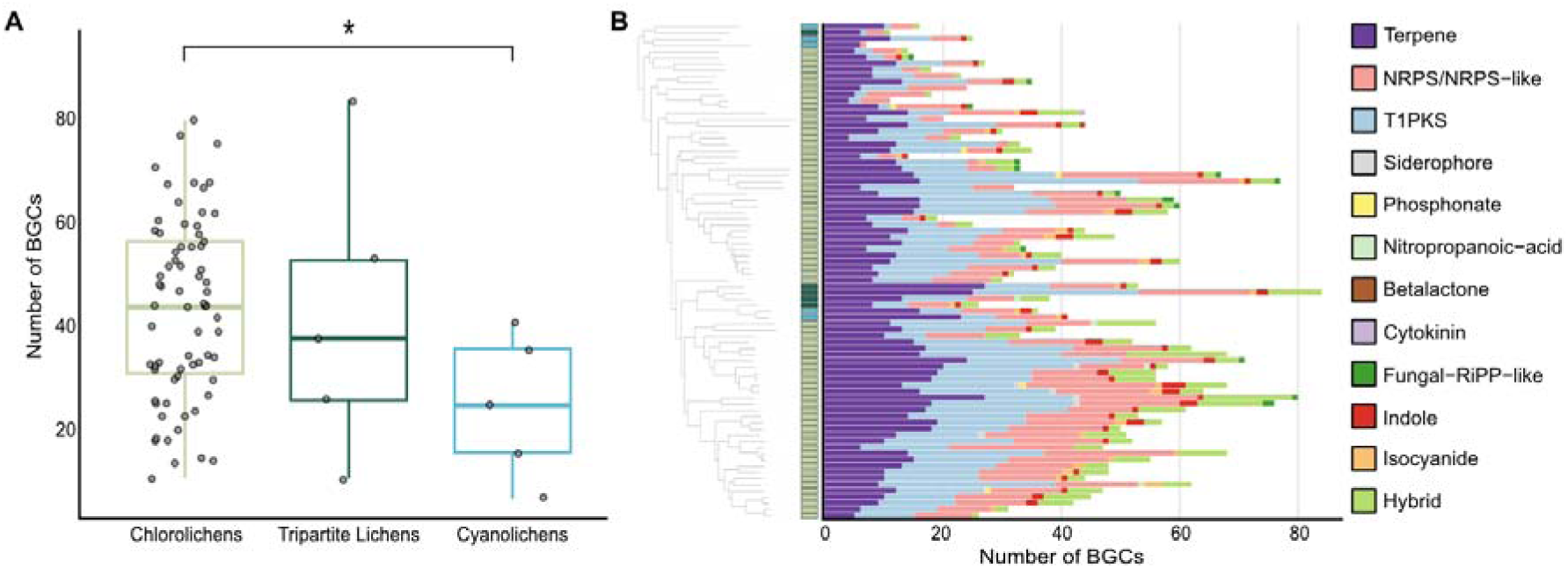
Abundance of biosynthetic gene clusters (BGCs) in lichen genomes. **A)** Box plot showing the number of BGCs encoded by mycobiont genomes grouped according to photobiont association. Each dot represents a single lichen species. Statistical significance is indicated by an asterisk (*; p < 0.05). The outlines of the box plots are colour-coded for bipartite cyanolichens (blue), bipartite chlorolichens (pale green), and tripartite lichens (dark green). **B)** Stacked bar plot showing the abundance and subclass composition of BGCs across 80 lichen species. Species are arranged according to the phylogenetic order shown in Fig. 1. Photobiont association types are indicated with bars: bipartite cyanolichens (blue), bipartite chlorolichens (pale green) and tripartite lichens (dark green). Colours of stacked bars indicate different BGC subclasses. Detailed counts for each BGC subclass are provided in Table S9.

BGCs are typically classified into subclasses based on the chemical nature of the products that the pathways produce (Dinglasan et al., 2025). Collectively, among the 80 species of lichen-forming fungi 11 distinct subclasses of BGCs were recorded using antiSMASH (Blin et al., 2025). BGCs containing genes from two or more distinct subclasses were annotated as hybrids. The 11 BGC subclasses included both broad-category mega enzyme clusters and clusters predicted to produce specific secondary metabolites. The broad-category BGCs comprised non-ribosomal peptide synthetase (NRPS) and type I polyketide synthase (T1PKS) gene clusters. The nine more specific BGC types identified target distinct structural or metabolite families, producing indoles, isocyanides, terpenes, cytokinins, betalactones, phosphonates, nitropropanoic acids, siderophores or fungal ribosomally synthesized and post-translationally modified peptide-like metabolites (fungal RiPP-like) (Fig. 6B).

We subsequently examined how differences in total BGC content could be attributed to variation across the 11 identified BGC subclasses and found that terpene BGCs were identified in all genomes and were encoded on average 12 times per species with the abundance ranging from as low as four in *Umbilicaria vellea* and *Candelina submexicana*, to as high as 27 in *Physcia stellaris* and *Pseudocyphellaria aurata*. The NRPS BGCs were also present in all genomes and were encoded on average ten times per genome, while the T1PKS BGCs were absent in only a single species (*Phylliscum demangeonii*) and reached up to 37 in *Thelotrema lepadinum,* with 14 per species on average (Table S2). In contrast, the remaining eight specific metabolite BGCs were rare, appearing less than once per species on average and never exceeding four clusters in any single genome (Fig. 6B; Table S2). When comparing BGC subclass abundance across photobiont association types, only T1PKS clusters differed significantly (Kruskal–Wallis test, adjusted p-value = 0.041). This difference was driven by a reduced abundance of T1PKS clusters in cyanolichens compared with chlorolichens (Dunn’s test, adjusted p-value = 0.005), whereas no significant differences were detected for any of the remaining BGC subclasses. The reduced abundance of T1PKS clusters therefore likely accounts for the lower total BGC content observed in cyanobacteria-associated lichens (Table S9). This finding is consistent with the generally lower secondary-metabolite diversity reported for cyanobacteria-associated lichens, which particularly lack polyketide-derived lichen substances such as depsides and depsidones (Diaz-Escandon et al., 2022). Together, these results indicate that BGC repertoires of lichenized fungi are dominated by terpene, NRPS and T1PKS clusters, whereas the remaining subclasses occur only sporadically across species.

## DISCUSSION

Our comparative analysis reveals extensive AMP repertoires across lichen-forming fungi, indicating a widespread genomic potential to produce antimicrobial proteins. This finding is consistent with previous observations that fungi across diverse lifestyles encode large AMP repertoires and supports the notion that protein-based microbial antagonism may represent an ancient component of fungal biology (Snelders et al., 2022; Mesny et al., 2026). It also fits with the broader observation that lichen-forming fungi possess extensive repertoires of lineage-specific secreted proteins and other interaction-related genes (Resl et al., 2022). A role for secreted AMPs in niche colonization has been particularly well demonstrated for fungal plant pathogens, where increasing evidence shows that AMPs are secreted as a virulence strategy to manipulate host-associated microbiota and suppress microbial antagonists to promote infection (Snelders *et al*., 2020; 2021; 2023; Kraege *et al*., 2026; Mesny *et al*., 2024; 2026). Notably, many of these effectors derive from ancient antimicrobial protein families that predate land plants and were subsequently co-opted for host colonization, with individual effectors showing distinct target specificities (Snelders *et al*., 2020, 2021; Mesny *et al*., 2026). However, the evolutionary forces shaping these repertoires in lichen-forming fungi have remained unclear. Here, we show that predicted AMP repertoire size is broadly similar among fungi associated with green algae, cyanobacteria, or both, whereas the composition of these repertoires varies extensively among species and populations. In contrast, CAZyme and BGC repertoires show substantially stronger associations with photobiont type. Together, these patterns point to distinct evolutionary constraints acting on different classes of fungal proteins involved in interactions with other organisms.

The broad similarity in AMP repertoire size across photobiont associations is particularly notable given not only the fundamentally different cell biology and physiology, but also the ecological differences among these symbioses (Dal Grande et al., 2018; Rolshausen et al., 2020; Rangel et al., 2021; Spribille et al., 2022). Lichen-forming fungi interact with photosynthetic partners while simultaneously encountering diverse bacterial and fungal communities, and antimicrobial molecules have long been proposed to contribute to microbial competition and chemical organization within lichen thalli (Calcott et al., 2018). Our data suggest that the genomic potential for protein-based antimicrobial activity may constitute a broadly conserved feature of lichen-forming fungi.

An important consideration in interpreting the apparent conservation of AMP repertoire size is the limited representation of cyanolichens and tripartite lichens in our dataset. Each of these association types is represented by only five species, substantially fewer than the chlorolichens, and moderate differences among association types may remain undetectable. Nevertheless, the overall pattern differs markedly from that observed for BGCs and CAZymes, for which association-type differences were readily apparent despite the same sampling framework. Larger comparative datasets, particularly with greater representation of cyanolichens and tripartite lichens, even though these are comparatively rare in nature (Honegger, 2008), will be required to determine whether more subtle association-specific differences exist.

The broad conservation of repertoire size masks substantial compositional turnover. Across the 80 fungal genomes, individual AMP orthogroups were highly variable in occurrence, and only a subset was broadly shared among species. Such a pattern could arise if antimicrobial capacity is subject to stabilizing selection at the level of overall repertoire size while individual AMP families remain free to be gained, lost, or replaced. The extensive turnover observed here is consistent with the dynamic evolution of fungal secreted proteins and other interaction-associated gene families documented across diverse fungal lineages (Wisecaver et al., 2014, Resl et al., 2022; Mesny et al., 2026).

The contrast with CAZyme and BGC repertoires further suggests that different classes of fungal proteins are subject to different evolutionary constraints. Previous comparative genomic analyses of lichen-forming fungi have similarly demonstrated substantial differences in carbohydrate-degradation and transport potential among fungal symbionts, emphasizing that the metabolic interface with the photobiont can differ considerably among lichenized fungi (Resl et al., 2022). Lichen-forming fungi also possess extensive biosynthetic potential, including numerous fungal BGCs, although the functions of many of these pathways remain unresolved (Betrand et al., 2018; Calcott et al., 2018; Singh et al., 2025). However, specialized metabolites are increasingly recognized as mediators of microbial interactions (Rangel et al., 2021; Mesny et al., 2023), whereas CAZymes mediate how fungi interact with host- and environment-derived carbohydrates and can include enzymes targeting microbial cell wall components (Wardman et al., 2022; Zheng et al., 2023). Recent comparative analyses across microbial and fungal lineages link expanded specialized-metabolite biosynthetic capacity with complex multicellularity and show that BGC-rich lineages are concomitantly enriched in CAZymes (Salamzade et al., 2026). In our dataset, genomes with larger BGC repertoires likewise tended to have larger CAZyme repertoires, consistent with this broader pattern. Overall, the evolution of fungal antimicrobial potential may thus involve multiple partially independent molecular repertoires. This decoupling may reflect differences in the costs, functional breadth, target ranges, and evolutionary dynamics of these molecular classes. AMP repertoires may provide a flexible baseline capacity for microbial antagonism, whereas enzymatic and secondary-metabolite repertoires may be more directly shaped by particular host or environmental contexts. The chemically rich nature of lichen symbioses and the contribution of multiple partners to their secondary-metabolite repertoire further support the view that chemical and protein-based antimicrobial strategies need not evolve in parallel (Calcott et al., 2018).

Our analysis of AMP sequences and predicted structures provides further evidence for substantial molecular diversification within this conserved quantitative framework. Most AMP clusters showed correspondence between sequence and predicted structural similarity, whereas highly divergent sequences did not generally resolve into common structural groups. Thus, within the sensitivity of our structural clustering approach, and in contrast to findings reported for various fungal plant pathogens (de Guillen et al., 2015; Derbyshire and Raffaele 2023; Seong & Krasileva, 2023), we find limited evidence for widespread structural convergence among highly divergent AMP sequences. Instead, AMP diversification in lichen-forming fungi appears to involve both lineage-specific sequence diversification within conserved structural classes and the presence of distinct AMP families. This observation is consistent with the broader principle that fungal antimicrobial proteins can retain structural or functional features despite substantial sequence divergence, although individual AMP families can also evolve rapidly (Mesny et al., 2026). More sensitive structural phylogenomics, together with experimentally determined structures, may reveal additional remote relationships that are not captured by the current clustering thresholds.

The population-level analysis of *Umbilicaria phaea* provides a complementary perspective on this repertoire turnover. AMP repertoires differed among populations and showed associations with elevation and photobiont composition, while photobiont turnover itself was evident across the geographic gradient (Dal Grande et al., 2018; Rolshausen et al., 2020, 2023). These observations are consistent with the possibility that local ecological conditions and photobiont associations contribute to AMP diversification (Grube et al., 2009, 2015; Aschenbrenner et al., 2014; Keller et al., 2026). However, these factors are not independent in the present sampling design: elevation, geography, photobiont composition, and potentially microbiome composition covary across populations. The observed associations therefore cannot distinguish photobiont-mediated selection from environmental effects, population structure, or other correlated ecological variables. Rather than demonstrating direct adaptation to photobiont identity, the population-level results indicate that AMP repertoires can vary substantially within a fungal species. Future studies integrating intra-species population genomics and broader sampling across populations will be important to identify signatures of adaptive diversification while minimizing the phylogenetic constraints inherent to the cross-species comparisons used here.

Phylogenetic structure represents a further consideration when interpreting associations between AMP repertoires and photobiont type. Cyanobacterial associations occur in phylogenetically distinct fungal lineages, consistent with the independent origins of lichenization and the repeated evolution of symbiotic associations in fungi (Miadlikowska et al., 2006; Spribille et al., 2022; Resl et al., 2022). Photobiont association is nevertheless not randomly distributed across the fungal phylogeny. Consequently, association-specific differences in individual AMP families may reflect photobiont-associated selection, shared evolutionary history, or both.

Finally, the present analysis addresses genomic potential rather than the realized antimicrobial activity of lichen-forming fungi. Transcriptomic and proteomic analyses under different symbiotic and environmental conditions will be required to determine which predicted AMPs are expressed, whether expression varies with photobiont identity or microbiome composition, and whether repertoire turnover translates into differences in antimicrobial activity. Functional characterization of representative AMP families will be particularly important for establishing whether sequence and structural diversification corresponds to differences in microbial targets or mechanisms of action. Such analyses will also be necessary to determine whether the similar sizes of predicted AMP repertoires reflect selection for a general antimicrobial function or instead emerges from the independent evolution of multiple lineage-specific proteins with different ecological roles, or a combination thereof.

Taken together, our results suggest that lichen-forming fungi maintain a broadly conserved genomic potential for protein-based antimicrobial activity while dynamically remodeling the constituent AMP families. The key feature is therefore not static conservation of individual antimicrobial proteins, but a decoupling between repertoire size and repertoire composition. The contrasting evolutionary patterns of AMPs, CAZymes, and BGCs further indicate that different components of fungal antimicrobial potential are shaped by distinct evolutionary forces. Lichen-forming fungi may thus maintain a conserved quantitative capacity for microbial antagonism while retaining substantial molecular flexibility in how that capacity is realized.

## MATERIALS AND METHODS

### Mycobiont genome collection

Publicly available genome assemblies of 78 lichen-forming fungi were gathered from the NCBI genome database and JGI MycoCosm. *Lasallia pustulata* was renamed to *Umbilicaria pustulata*. Two additional genomes sequenced at the Institute for Biological Data Science of the Heinrich-Heine-University Düsseldorf were added resulting in a total of 80 genome assemblies (Joisten-Rosenthal et al., 2026; Fig. 1; Table S1).

### Phylogenomic analysis

OrthoFinder v2.5.5 (Emms et al., 2019) was utilized to generate a species tree based on proteomes extracted from the genomes, using the implemented method STAG, which was visualized with iTOL v7 (Letunic and Bork, 2024).

### Genome annotation

All genomes were analysed for genome completeness using BUSCO v5.8.0 with lineage dataset ascomycota_odb10 (Manni et al., 2021). To ensure comparability, all genome assemblies were re-annotated using BRAKER v3.0.8 (Gabriel et al., 2024) (with option ‘--fungus’ and other parameters set to default) which predicts protein coding gene structures by integrating GeneMark-ETP (from the GeneMark-ES Suite v4.71; Brůna et al., 2024; Ter-Hovhannisyan et al., 2008) and AUGUSTUS 3.5.0 (Stanke et al., 2004). Secondary metabolite biosynthesis gene clusters (BGCs) were annotated with the antiSMASH 8.0 web server (Blin et al., 2025) set to detection strictness “relaxed”. SignalP 6.0 (Teufel et al., 2022) was run on the BRAKER-annotated protein sequences to predict signal peptides for secretion. All proteins predicted to carry a signal peptide were considered as “secretome”. Carbohydrate-active enzymes (CAZymes) were annotated with dbCAN v3 (Zheng et al., 2023).

### Structure prediction and AMP protein clustering

To predict the antimicrobial properties of the secretome, we first predicted structures of the secreted proteins, with an upper limit of 800 amino acids, using ESMFold v1.0.3 (Lin et al., 2023). As only few carbohydrate-active enzymes (CAZymes) have been included in the AMAPEC training set, and thus antimicrobial activity predictions for them are less robust than for other types of secreted proteins, CAZymes were omitted from the structure prediction. The ESMFold-predicted structures were used to identify putative antimicrobial proteins using AMAPEC v1.0 (Mesny et al., 2026). OrthoFinder was re-run on the secretome to assign proteins into orthogroups. To assess structural similarities, in addition to the sequence similarity, we used Foldseek v9.427df8a (Van Kempen et al., 2024) easy-cluster to combine structurally similar proteins using a TM-align cut-off ≥ 0.5.

### Intraspecific AMP variation across elevations

Pool-Seq data from 11 *Umbilicaria phaea* populations sampled along two elevational gradients in California were used to assess intraspecific AMP variation. AMP catalogues were generated using AMAPEC v1.0 (Mesny et al., 2026) for two available *U. phaea* reference genomes. The resulting 325 AMP candidates were screened against the Pool-Seq data from all 11 populations to assess their presence–absence patterns across populations.

### Statistics

Statistical analyses were performed in R. Differences among photobiont association types were assessed using Kruskal–Wallis tests followed by Dunn’s tests for pairwise comparisons.

Two-group comparisons were performed using Student’s t-tests or Wilcoxon tests depending on the distribution of the data, as assessed using Shapiro tests. Associations between AMP repertoire size, genome size, secreted protein number and BGC abundance were assessed using Spearman rank correlations. AMP orthogroup and structural-cluster occurrence patterns were compared using Fisher’s exact tests. P values were adjusted for multiple testing using the Benjamini–Hochberg method where applicable.

## ACKNOWLEDGEMENTS

This study was supported by Deutsche Forschungsgemeinschaft (DFG) within the framework of CRC1535 project no. 458090666. B.P.H.J.T. acknowledges funding by the Alexander von Humboldt Foundation in the framework of an Alexander von Humboldt Professorship endowed by the German Federal Ministry of Education and Research. B.P.H.J.T. and B.U. are furthermore supported by DFG under Germany’s Excellence Strategy – EXC 2048/1 – Project ID: 390686111. The authors thank Fantin Mesny, Anton Kraege and Vittorio Tracanna for their support and guidance with computational approaches.

## AUTHOR CONTRIBUTIONS

C.K., H.R., and B.P.H.J.T. conceived the project. C.K., E.L.Y.W., I.S. and B.P.H.J.T. designed the analyses. C.K., E.L.Y.W., and A.v.W. analysed the data. V.J., and B.U. provided genomes. C.K. and B.P.H.J.T. wrote the manuscript. All authors read and approved the final manuscript.

