## Supplementary Figures for "Decoupled evolution of antimicrobial repertoires in lichen-forming fungi"

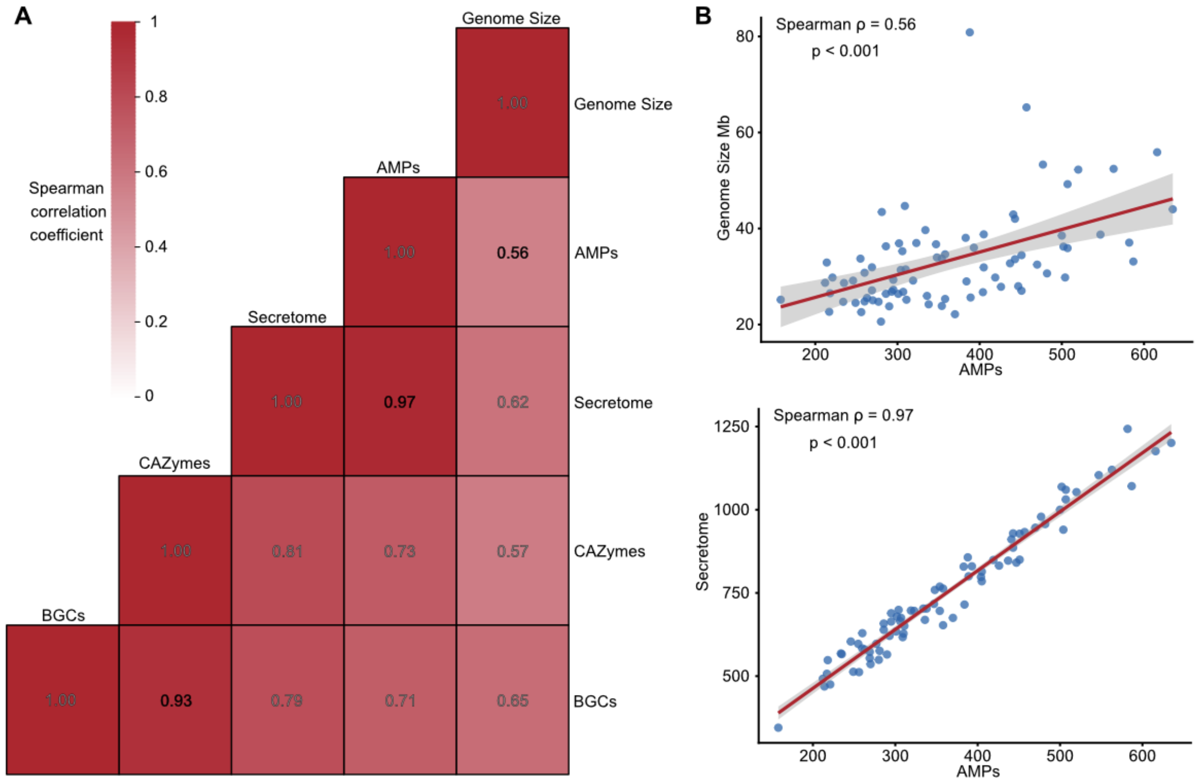


**Figure S1. Spearman correlations among genomic and predicted antimicrobial molecule repertoires in lichen-forming fungi. A)** Heatmap showing pairwise Spearman correlations between genome size, number of predicted secreted proteins, antimicrobial proteins (AMPs), carbohydrate-active enzymes (CAZymes) and biosynthetic gene clusters (BGCs) across the 80 lichen-forming fungal species. Correlation coefficients are indicated by colour, with zero correlation shown in white and increasing positive correlations shown in dark red Correlation coefficients corresponding to values reported in the Results are shown in black, while all other coefficients are shown in grey. **B)** Scatter plots showing the relationships between AMP number and genome size (Spearman ρ = 0.56) and between AMP number and number of predicted secreted proteins (Spearman ρ = 0.97).

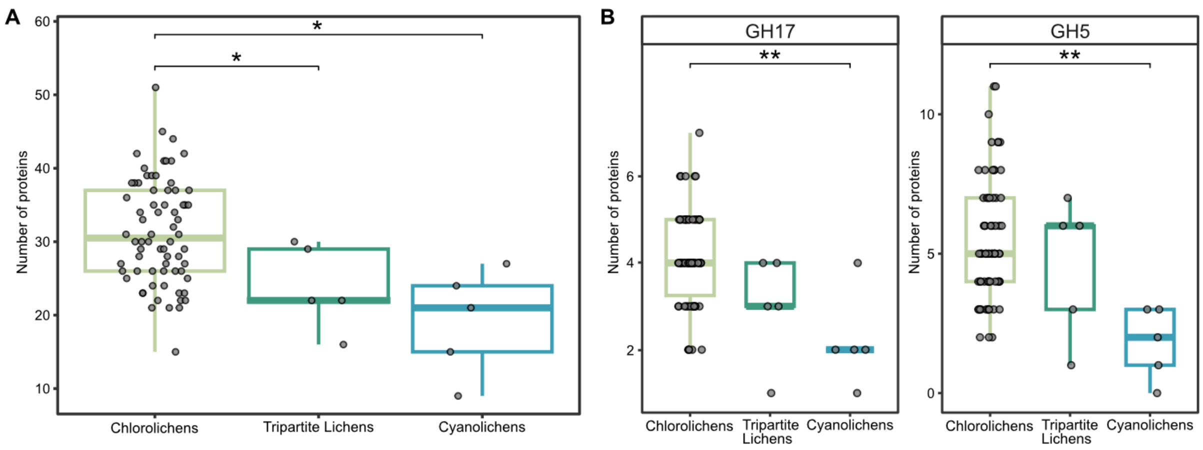


**Figure S2. Comparison of cell-wall-associated CAZyme families across lichen-forming fungi with different photobiont associations. A)** Box plots showing the combined abundance of CAZyme families with reported activity against microbial or algal cell-wall components, including families targeting bacterial peptidoglycan, fungal and algal chitin, and fungal and algal β-glucans, as well as AA11. Bipartite chlorolichens encoded significantly more of these cell-wall-associated CAZymes than bipartite cyanolichens or tripartite lichens (pairwise Wilcoxon test, adjusted *p-value = 0.011 and *p-value = 0.049). The outlines of the box plots are colour-coded for bipartite cyanolichens (blue), bipartite chlorolichens (pale green), and tripartite lichens (dark green). **B)** Box plots showing the abundance of individual cell-wall-associated CAZyme families. GH17 and GH5 were significantly more abundant in bipartite chlorolichens than in bipartite cyanolichens (pairwise Wilcoxon test, adjusted **p-value = 0.0087 and 0.0022, respectively). The outlines of the box plots are colour-coded for bipartite cyanolichens (blue), bipartite chlorolichens (pale green), and tripartite lichens (dark green).
